# CompAIRR: ultra-fast comparison of adaptive immune receptor repertoires by exact and approximate sequence matching

**DOI:** 10.1101/2021.10.30.466600

**Authors:** Torbjørn Rognes, Lonneke Scheffer, Victor Greiff, Geir Kjetil Sandve

## Abstract

Adaptive immune receptor (AIR) repertoires (AIRRs) record past immune encounters with exquisite specificity. Therefore, identifying identical or similar AIR sequences across individuals is a key step in AIRR analysis for revealing convergent immune response patterns that may be exploited for diagnostics and therapy. Existing methods for quantifying AIRR overlap do not scale with increasing dataset numbers and sizes. To address this limitation, we developed CompAIRR, which enables ultra-fast computation of AIRR overlap, based on either exact or approximate sequence matching. CompAIRR improves computational speed 1000-fold relative to the state of the art and uses only one-third of the memory: on the same machine, the exact pairwise AIRR overlap of 10^4^ AIRRs with 10^5^ sequences is found in ∼17 minutes, while the fastest alternative tool requires 10 days. CompAIRR has been integrated with the machine learning ecosystem immuneML to speed up various commonly used AIRR-based machine learning applications.

**Availability and implementation:** CompAIRR code and documentation are available at https://github.com/uio-bmi/compairr. Docker images are available at https://hub.docker.com/r/torognes/compairr. The scripts used for benchmarking and creating figures, and all raw data, may be found at https://github.com/uio-bmi/compairr-benchmarking.

## Introduction

Adaptive immune receptor (AIR) repertoires (AIRRs) record past immune encounters. High-throughput sequencing now enables millions of AIR sequences to be determined at a cost that facilitates adaptive immunity-based association studies on large patient cohorts^1,2^. It has been previously shown that shared immune states give rise to identical or similar AIR sequences across individuals, enabling the use of AIRR-seq for diagnostics and therapeutic research^3,4^. Computation of cross-individual AIRR intersections, i.e., the number of matching AIR sequences across AIRRs, is thus a foundational computational task performed in nearly all AIRR analyses. However, since the number of pairwise AIRR comparisons grows quadratically with the number of AIRRs considered, where each pairwise AIRR comparison typically involves millions of individual AIRs, computational efficiency is crucial for performing AIR sequence matching at scale.

We here present CompAIRR, a tool that allows to compute AIRR intersections up to 1000-fold faster than current implementations^5–7^. In contrast to existing tools, CompAIRR supports both exact and approximate sequence matching between AIRs when determining AIRR overlap. The implementation is available as a stand-alone command-line tool, and it is integrated with the machine learning ecosystem immuneML8 to accelerate the computation of AIRR similarity matrices, and to accelerate an AIRR-based immune state classifier by Emerson et al.^1^ (Supplementary Figure 1).

## CompAIRR description

CompAIRR is based on a sequence comparison strategy developed for the nucleotide sequence clustering tool Swarm^10^. A Bloom filter^11^ and a hash table are used to quickly look up similar AIR sequences across AIRR sets. For each AIR sequence (nucleotide or amino acid), a 64-bit hash value is generated using a Zobrist hash function^12^, a form of tabulation hashing that can be computed efficiently and updated incrementally. When approximate matching is enabled, the hashes of all possible variants of a query sequence (with 1–2 substitutions or indels) are also generated. Matches are optionally restricted by V and J gene. Multi-threading may be enabled to further speed up comparisons (see figure 1d). For the comparison of *n* AIRRs, CompAIRR produces an *n*x*n* matrix where each cell contains the sum of matching AIR frequencies with flexible summary statistics (product, min, max, mean or ratio of the two compared AIR frequencies). Alternatively, CompAIRR can query *n* AIRRs against *m* reference AIRs and produce an *n*x*m* sequence presence table. While AIR matching is only supported at the single chain level, two *n*x*m* sequence presence tables for complementary (paired) AIR chains (single-cell data) can easily be merged. CompAIRR can optionally output the list of (approximately) matching AIRs as an AIRR-compliant TSV file, and adheres to the AIRR standard for software tools ^13^.

**Figure 1.**
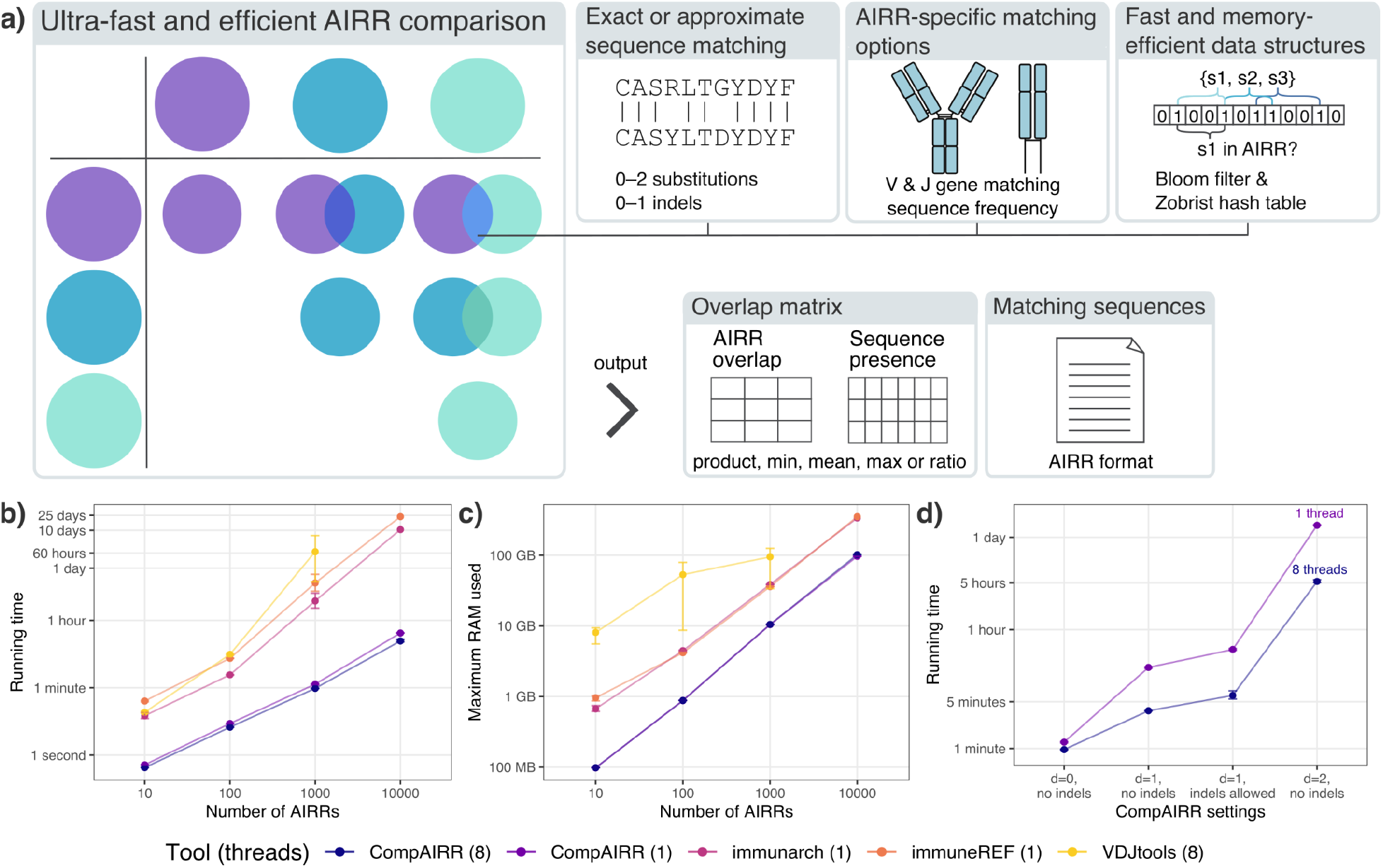
Overview of CompAIRR features and performance. **(a)** CompAIRR has configurable AIR matching criteria and output formats. **(b)** CompAIRR calculates pairwise AIRR overlap up to 1000-fold faster than currently available tools. **(c)** The maximum RAM usage of CompAIRR is below one-third of the most memory-efficient alternative. **(d)** The CompAIRR running time increases when allowing more AIR sequence mismatches, but multithreading helps reduce this running time. **(b-d)** Data shown are mean with error bars showing min/max values across three replicate runs. For the largest dataset, only CompAIRR was run three times, and VDJtools failed to run due to memory limitations. Unless otherwise specified, datasets consist of 1000 AIRRs containing 10^5^ OLGA^9^-generated sequences (default human IgH CDR3 model).

## CompAIRR performance benchmarking

CompAIRR (1.3.1) was benchmarked against VDJtools (1.2.1)^5^, immunarch (0.6.5)^6^ and immuneREF (0.5.0)^7^ by calculating the pairwise AIRR overlap of datasets ranging from 10 to 10^4^ AIRRs. Each AIRR consisted of 10^5^ amino acid AIR sequences generated using OLGA^9^ with the default human IgH CDR3 model. Figures 1b and 1c respectively show the running time and maximum RAM usage of each tool. CompAIRR is consistently faster, particularly for large datasets: with 10^4^ AIRRs of 10^5^ sequences, CompAIRR ran in 17 minutes while immunarch took 10 days, immuneREF took 23 days and VDJtools failed to complete due to memory constraints. The computational complexity appears to have been reduced from approximately quadratic to almost linear. Furthermore, the maximum RAM usage of CompAIRR is below one-third of that of competing tools. The running time and memory usage as a function of the AIRR size (10^4^–10^6^ sequences) is shown in Supplementary Figure 2.

Additionally, Figure 1d shows how the CompAIRR running time is affected by approximate sequence matching, which is not at all supported by the existing tools. The benefit of multi-threading becomes more apparent when the degree of sequence mismatching is increased, since with exact matching the running time is dominated by disk access (Supplementary Figure 3).

## Conclusion

Identification of shared AIRs across AIRRs from different individuals is a core computational task in AIRR analysis. We have here presented CompAIRR, which calculates AIRR overlap up to 1000-fold faster while its peak memory usage is below one third compared to currently available tools. We validated that CompAIRR easily scales to datasets of 10^4^ AIRRs of 10^5^ sequences each, which surpass the largest available experimental datasets^2,14^. Furthermore, a novel feature of CompAIRR is efficient identification of *approximately* matching AIR sequences across AIRRs or to reference databases, which may be a biologically meaningful way to increase the number of matches between AIRRs when the exact overlap is low (Supplementary Figure 4).

Complementary to sequence-level clustering tools ClusTCR^15^ and GIANA^16^, or comparison of AIRR subsets^17^, CompAIRR can be used for ultrafast similarity-based comparison of *complete* AIRRs. Due to flexible specification of summary statistics and output, CompAIRR is easily integrated with any tool capable of reading in either (i) a pairwise distance matrix containing cross-AIRR matches, (ii) a matrix showing individual AIR presence in one or more AIRRs, or (iii) an AIRR-compliant TSV file containing (approximately) matching AIRs between AIRRs. This allows accelerating a variety of analyses where AIRR comparison is a core computational component, including AIRR similarity and clustering^5,18^, graph analysis^19–21^, and immune state classification^1^.

## Acknowledgements

We are grateful to Milena Pavlović for her assistance. We are also grateful for access to computational resources provided by the Norwegian Research and Education Cloud (NREC) and UNINETT Sigma2 (project NN9383K).

## Supplementary figures

**Supplementary Figure 1.**
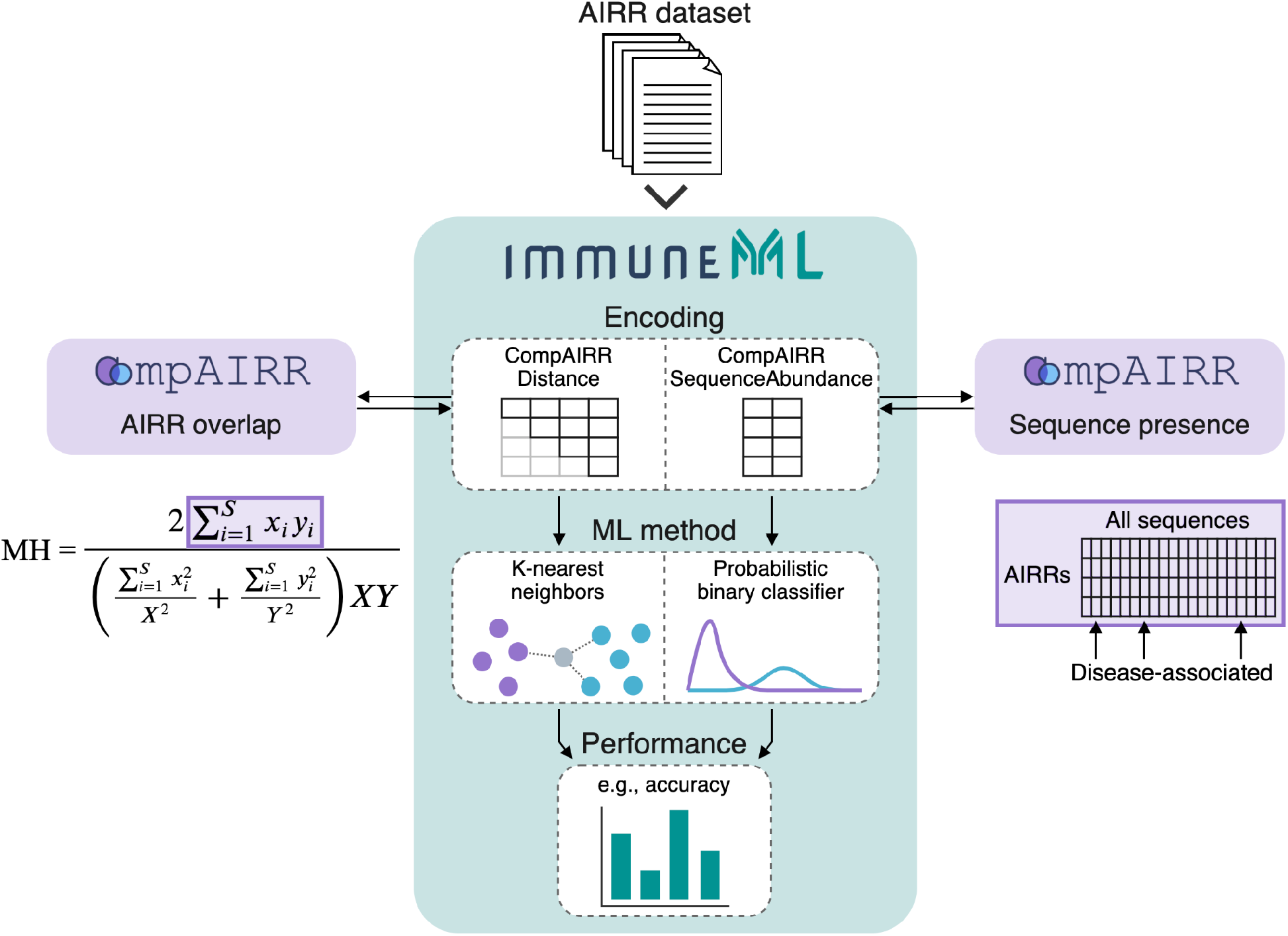
CompAIRR is integrated with immuneML^8^ to speed up core calculations of two different encodings. Firstly, CompAIRR accelerates the calculation of a Morisita-Horn (MH) distance matrix by calculating the sum of the products of overlapping AIR frequencies between AIRRs (left, highlighted in purple). This distance matrix can subsequently be used in combination with a k-nearest neighbors classifier. Secondly, calculating a sequence presence matrix with CompAIRR (right, highlighted in purple) speeds up the method described by Emerson et al.^1^, which was replicated in the immuneML ecosystem by a sequence abundance encoding (AIRR size and number of disease-associated AIRs per AIRR) and a probabilistic binary classifier.

**Supplementary Figure 2.**
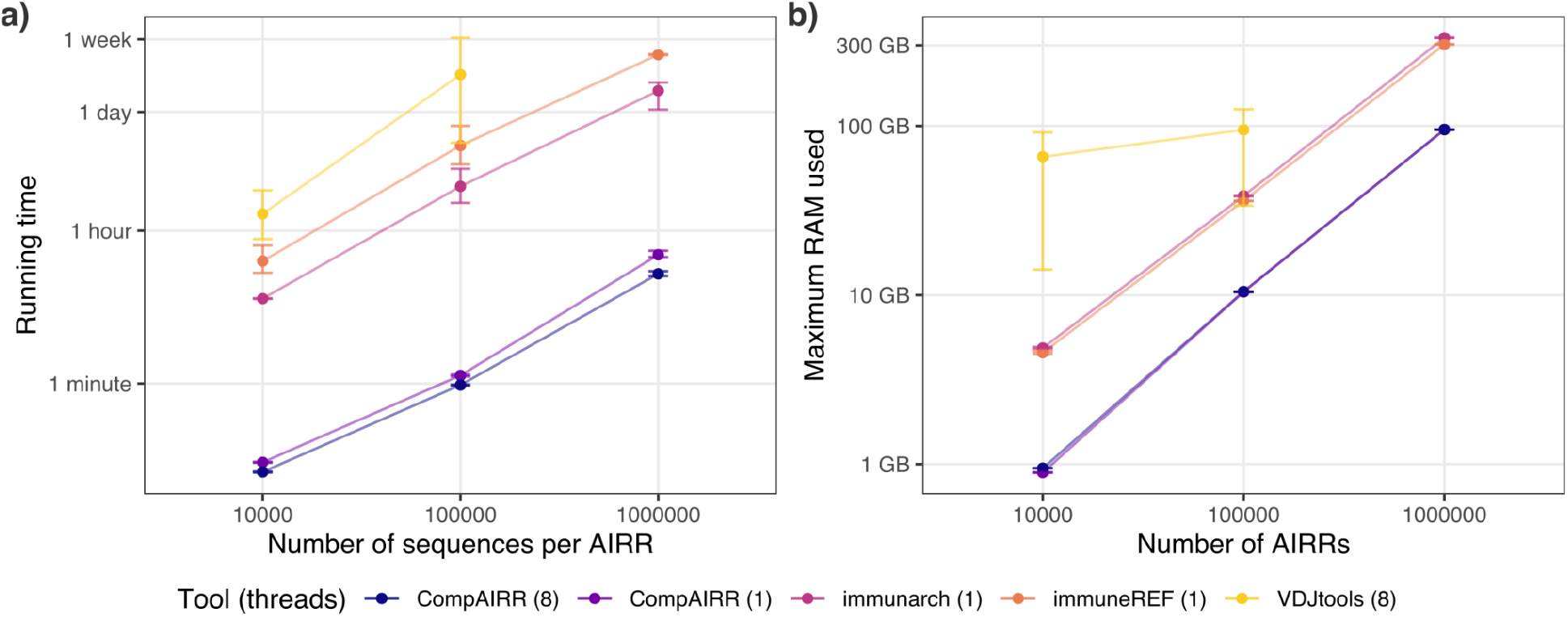
Benchmarking of (a) running time and (b) maximum RAM usage as a function of the number of sequences per AIRR. Data shown are mean with error bars showing min/max values across three replicate runs. Datasets consist of 1000 AIRRs (OLGA^9^-generated human IgH CDR3 sequences). VDJtools failed to run on the dataset of 10^6^ AIRRs due to memory limitations.

**Supplementary Figure 3.**
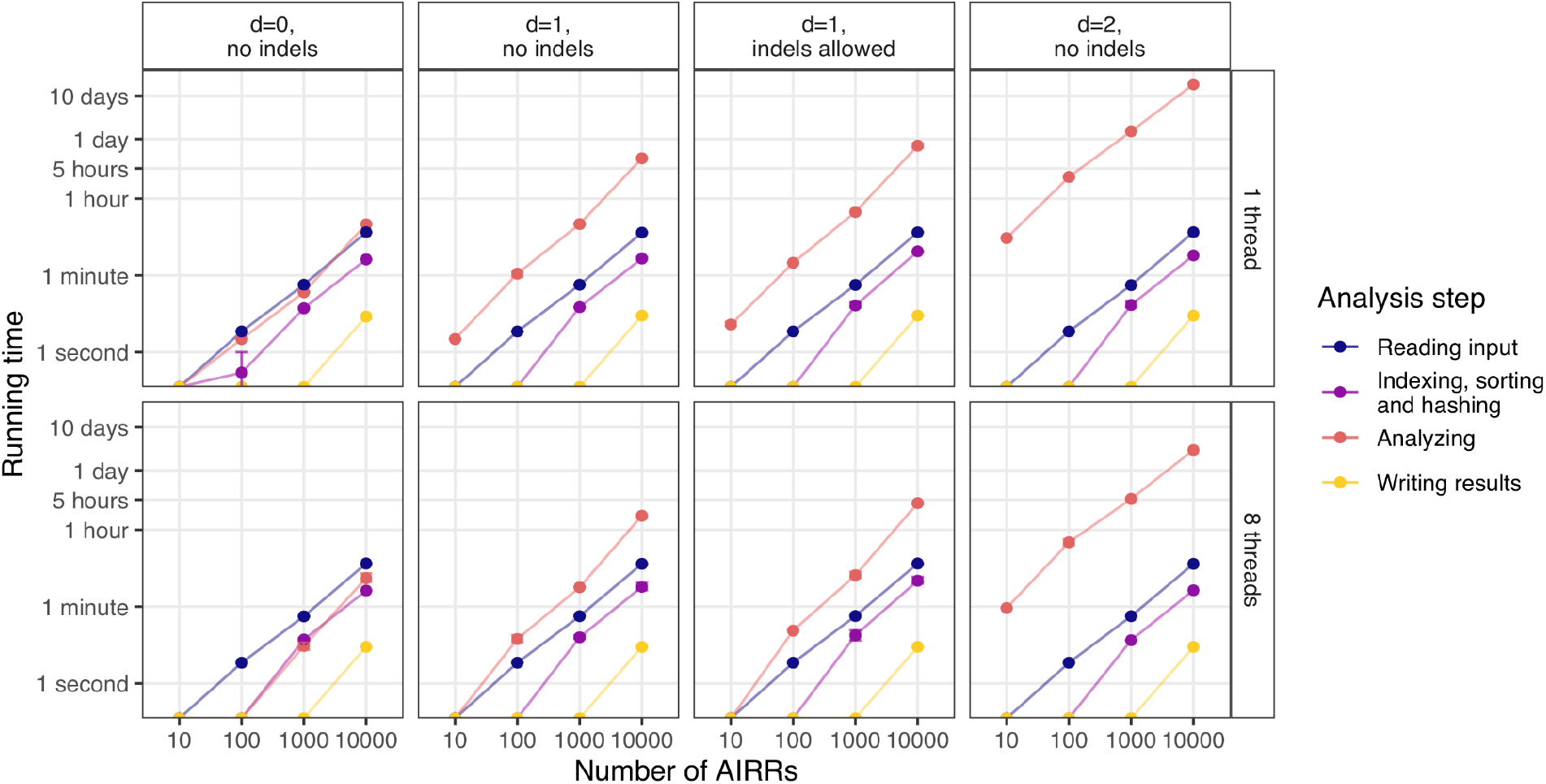
Running time of CompAIRR analysis steps. Reading input and analyzing the AIRR overlap are the most computationally intensive steps. When exact sequence matching is used, reading input data takes a relatively large amount of the total running time. As a result, the benefit of multi-threading is more apparent when non-exact matching is used.

**Supplementary Figure 4.**
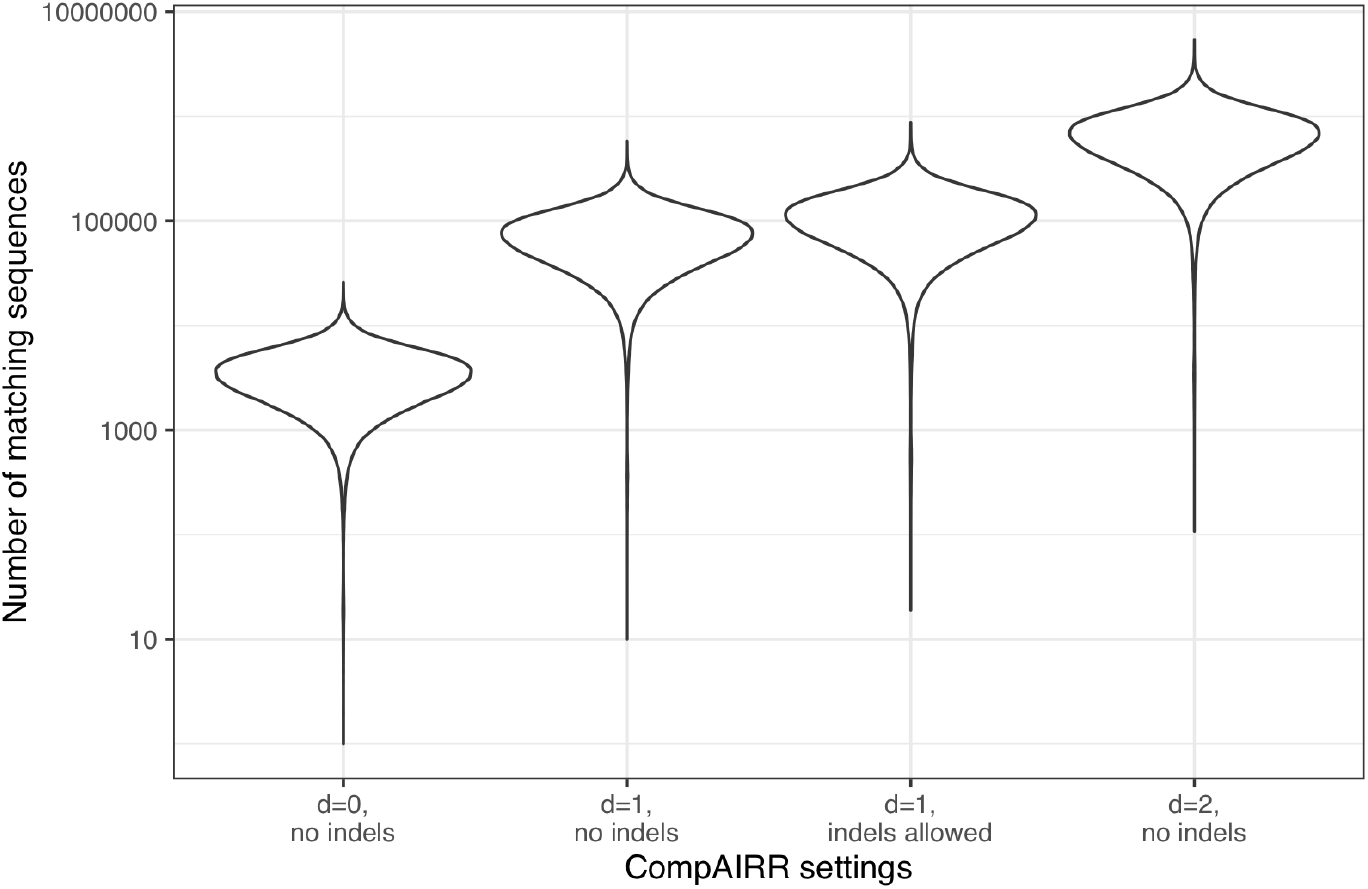
Number of matches with approximate sequence matching settings. The number of matching TCRβ AIR sequences between each combination of 710 AIRRs published by Emerson et al.^1^ increases when less stringent sequence matching criteria are used. When no or few exact overlapping sequences are found across AIRRs, highly similar sequences may still be present.

